# Unveiling Vaginal Fibrosis: A Novel Murine Model Using Bleomycin and Epithelial Disruption

**DOI:** 10.1101/2023.12.18.572175

**Authors:** Jennifer M McCracken, Gisele A Calderon, Lishore A Kumar, Swathi Balaji, Felipe Rivas, Dorothea Erxleben, Adam Hall, Julie CE Hakim

**Affiliations:** Baylor College of Medicine, Department of Obstetrics and Gynecology, Division of Pediatric and Adolescent Gynecology, Houston, Texas, USA; Texas Children’s Hospital, Department of Surgery, Division of Pediatric Surgery, Houston, Texas, USA; Wake Forest University School of Medicine, School of Biomedical Engineering and Sciences

**Keywords:** vagina, wound healing, hyaluronan, collagen, fibrosis, epithelial disruption

## Abstract

**Objective:** Develop, validate, and characterize a fibrotic murine vaginal wound healing model using bleomycin instillations and epithelial disruption.

**Approach:** We tested the effect of repeated bleomycin instillations with mucosal layer disruption on induction of vaginal fibrosis. Tissue samples collected at various time points were analyzed for fibrosis-related gene expression changes and collagen content.

**Results:** Low (1.5U/kg) and high-dose (2.5U/kg) bleomycin instillations alone did not induce fibrosis, but when high-dose bleomycin was combined with epithelial disruption, increased pro-fibrotic gene expression and trichrome staining were observed. To evaluate spatial and temporal changes in the ECM structure and gene expression, tissue samples were collected at 1 day, 3 weeks, and 6 weeks after bleomycin and epithelial disruption. Data analyses revealed a significant decrease in matrix metabolizing genes and an increase in pro-fibrotic genes and inhibitors of matrix metabolizing genes in the bleomycin plus epithelial disruption group at 3 weeks. Elevated levels of the profibrotic genes *Acta2*, *Col1a1*, and *Col3a* were exclusively detected in this group at 3 weeks, and trichrome staining confirmed increased collagen content after 3 weeks. Hydroxyproline levels showed a tendency towards elevation at 3 weeks (p=0.12) and 6 weeks (p=0.14), indicating fibrosis manifestation at 3 weeks and resolution by 6 weeks post-instillation and epithelial disruption.

**Innovation:** We combined bleomycin instillations with epithelial disruption to induce fibrosis and understand the mechanisms of the vaginal repair process.

**Conclusions:** Epithelial disruption combined with bleomycin induces murine vaginal fibrosis within three weeks, characterized by increased collagen synthesis. Remarkably, the vaginal tissue fully recovers within six weeks, elucidating the regenerative capacity of the vagina.

## INTRODUCTION

The vagina can sustain injury and immense tensional forces, such as those experienced during childbirth via vaginal delivery and still has the remarkable potential for regenerative healing.^1^ Under different wound healing conditions, such as reconstruction to correct congenital birth defects or after radiation exposure, the vagina has the potential to develop fibrosis and scar formation, causing significant pain and psychologic burden.^2^ However, the specific cellular and molecular mechanisms that determine whether wound repair follows a regenerative or fibrotic pathway remain unknown. Considering the high burden of fibrotic vaginal wound complications, seen in up to 73% of post-surgical vaginal wound beds^3^, research in this area is vital for improving global women’s health outcomes. A considerable number of girls and women who experience vaginal fibrosis, regardless of the underlying cause, are likely to develop some degree of vaginal stenosis or lumen narrowing. This may necessitate surgical intervention or ongoing vaginal dilatation for the remainder of their lives.^4^ Therefore, to improve the quality of life for patients affected by vaginal fibrosis, we seek to identify the mechanisms during the vaginal repair process that contribute to uncontrolled, disordered healing in order to develop techniques and therapies aimed at prevention and early targeted intervention.

We previously established a surgical murine vaginal wound model, using a 1 mm full thickness punch biopsy which healed regeneratively, or without fibrosis, 72 hours after injury.^5^ Given our previous work and the imperative need to create a vaginal injury model leading to fibrotic outcomes, our primary objectives of this project were to develop, validate, and characterize a model of murine vaginal fibrosis. In this work, we use repeated bleomycin vaginal instillations combined with traumatic disruption of the mucosal layer using bristle curettage to induce murine vaginal fibrosis.

Bleomycin is a clinical chemotherapeutic agent and bleomycin instillation is widely used to induce experimental fibrosis in multiple organ systems such as skin and lung^6,7^ where it has been shown to increase collagen deposition and hydroxyproline levels, cardinal features of organ scarring.^7,8^ In the case of laryngotracheal stenosis induction, bleomycin coated wire brushes were used to cause both physical and chemical injury which in turn induced scar formation.^9^ Though this technique has not been previously used to induce vaginal fibrosis, the vaginal mucosa bears a striking resemblance to other mucosal tissues that have been studied previously, suggesting that this may be a viable method for inducing vaginal fibrosis.

### Innovation

Vaginal fibrosis is a condition affecting millions of women worldwide that has received limited attention compared to fibrosis in other organ systems. Currently, *in vivo* fibrosis models exist for various organs, but none specifically focus on the vagina, hindering our understanding of its progression and healing mechanisms. We address this gap by creating a new murine model of vaginal fibrosis by combining bleomycin instillation with epithelial disruption to effectively trigger fibrosis. Our findings align with laryngotracheal models that adopted similar technique of fibrosis induction, and shed light on the mechanisms underlying vaginal fibrosis progression and resolution. These insights have the potential to enhance our understanding of this condition and guide the development of preventive and targeted treatment approaches to attenuate vaginal fibrosis.

### Clinical Problem Addressed

Collectively, this work will enable us to better understand the pathogenesis of disordered vaginal wound repair and fibrosis with the overarching goal of improving patient treatment and quality of life. Our goal was to develop a reproducible vaginal fibrosis model in mice to fundamentally understand potential healing mechanisms and what differs between regenerative healing and fibrotic healing. We hope to ultimately develop treatments and prevention options for the women and adolescents who undergo pelvic radiation and/or surgery who often face lifelong sequelae from resulting vaginal fibrosis with few treatment options.

## MATERIALS AND METHODS

### Animal Care and Use

C57BL/6J female mice between 6-7 weeks old were purchased from the Center for Comparative Medicine at Baylor College of Medicine (BCM) and were socially housed under a 12h/12h light/dark cycle with free access to standard food and water. All animal research was approved by the Institutional Animal Care and Use Committee (IACUC) of BCM, AN-7475, and conducted in accordance with the NIH Guide for the Care and Use of Laboratory Animals.

### Vaginal Injury

Bleomycin sulfate (B8416-15U, Sigma Aldrich, St. Louis, MO) was reconstituted with sterile normal saline. Mice were weighed prior to each exposure and bleomycin doses were adjusted according to animal weight to achieve weight-corrected volumes of either low dose (1.5U/kg) or high dose (2.5U/kg) bleomycin vaginal instillations. Normal saline intravaginal instillations were used as control. Three experimental conditions included: low dose bleomycin, high dose bleomycin, and epithelial disruption followed by high dose bleomycin. Epithelial disruption was done by inserting a wire brush (0.079” diameter) intravaginally until it met resistance and rotating clockwise 6 times. Mice were sedated using isoflurane at a rate of 0.5L/min. Eloxiject (Meloxicam, 5mg/mL, Henry Schein NDC 11695-6925-1) was given to each mouse at a dose of 5mg/kg prior to sedation. Normal saline or bleomycin was instilled intravaginally (with or without prior epithelial disruption) using a p100 pipette tip. Mice were kept sedated, with tail held suspended for 5 minutes to allow absorption, followed by applying an aquaphor healing ointment “plug” across vaginal opening. They were then transferred to clean caging and allowed to recover. This was repeated 5 times over the course of 10 days. Mice were euthanized either 1 day, 3 weeks, or 6 weeks after the last installation. Vaginal tissue was then collected and either placed in 10% formalin (n=7-8) or snap frozen for future analysis (n=7-8 for gene expression, n=3 for HA analysis).

### Histology and immunohistochemistry

Tissue placed in formalin at time of collection was dehydrated through a series of graded ethanols and embedded in paraffin (FFPE). Sections in the size of 5µm were then made for staining.

To evaluate tissue fibrosis and collagen content, sections were stained using Gomori’s Trichrome stain (Leica, Buffalo Grove, IL) according to manufacturer’s directions using Leica Autostainer. Images were taken at 20X using Leica DMi8 microscope with a Leica DFC4500 camera and Leica Application Suite X (LAS-X). ImageJ was used to determine the total area of submucosal tissue and amount of collagen content (blue staining) in the area using the color masking plug in as described previously ^8, 9^. Settings were: Hue: 124-194, Saturation: 0-255, Brightness: 0-255. Data is reported as % positive area, and 3 images per mouse were averaged.

Immunohistochemistry was used to localize total leukocytes and macrophages in the vaginal submucosa as described previously.^3^ Briefly, FFPE sections were rehydrated, and the PT Link antigen retrieval system (Agilent, Santa Clara, CA) and low target retrieval solution (Agilent) were used. The Dako Autostiner Link 48 (Agilent) and appropriate Agilent reagents were used for blocking (S2003), antibody dilution (S0809), secondary antibody (K4009), and 3,3’-diaminobenzidine to visualize (DAB, K3468, Agilent). Hematoxylin (K8008) was used as a counterstain. CD45 (ab10558, 1:2000, AbCam) and F480 (MF48000, Thermo Fisher) primary antibodies were used. A Leica DMi8 microscope with Leica DFC4500 camera and LAS-X software was used to take 20X images (n=3/mouse). ImageJ was used to determine the submucosal area in each image and to count the number of positive cells to calculate the number of positive cells/submucosal area.

### Gene Expression

Vaginal tissue was flash frozen in liquid nitrogen at time of collection and stored at 80°C until RNA was isolated using a PureLink RNA Minikit (ThermoFisher) with an added DNA elimination step according to manufacturer’s directions.

RNA was pooled so that there were equal amounts of RNA in each group per time point and each mouse in the group had equal contributions to the pooled sample. RNA was reverse transcribed using RT^2^ First Strand according to manufacturer’s instructions (Qiagen, Germantown, MD). RT^2^ Profiler PCR Array for Mouse Fibrosis (PAMM-120Z, Qiagen) and Qiagen GeneGlobe Analysis was used to evaluate fibrosis between normal saline and experimental groups at each time point.

Standard qPCR was used to validate a selection of genes from the array. In brief, 220ng of RNA was reverse transcribed using High-Capacity RNA-to-cDNA kit (Fisher Scientific). Power SYBR green and Bio-Rad CFX thermocycler was used. Samples were done in duplicate, and data was analyzed using the △△Ct method and expressed as fold change over saline control at each time point. BioRad PrimePCR SYBR green assay (Cat 10025636) was used. Additional primers used are listed in Table 1.

**Table 1.**
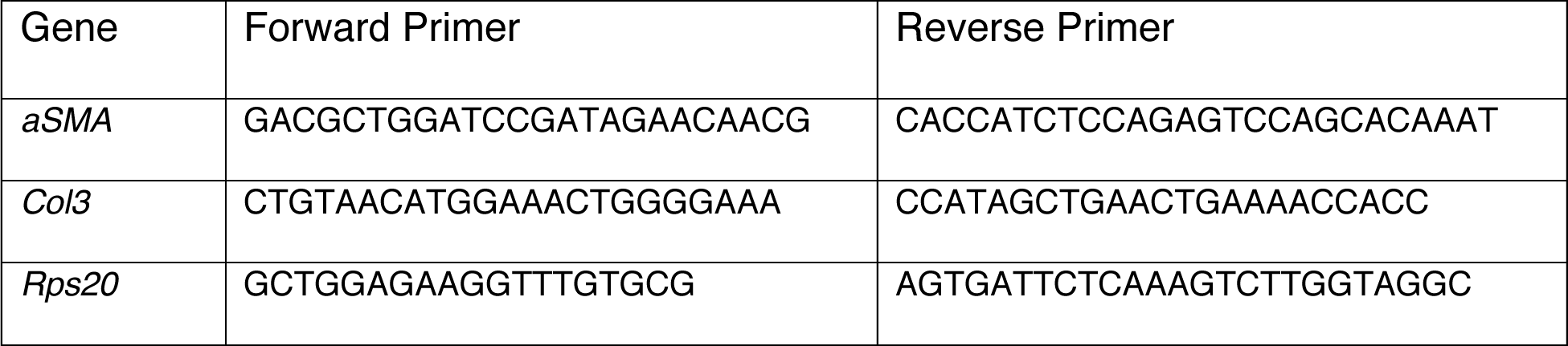

### Hydroxyproline

Vaginal hydroxyproline content was used as a surrogate marker for collagen.^10^ The hydroxyproline assay kit (Sigma) was used according to manufacturer’s direction. Briefly, tissue was homogenized in 100μL of water, combined with 100μL of 12N HCl and heated for 3 hours at 120°C. The tissue was vortexed every 30 minutes and, after heating, spun at 10,000g for 3 min. A 96 well plate was loaded with provided hydroxyproline standard and samples and allowed to dry completely in a 60°C oven. For each well, 6μL of Chloramine T and 94μL of Oxidation buffer was mixed, added, and allowed to incubate for 5 minutes at room temperature. DMAB concentrate was diluted 1:2 with Perchloric Acid/Isopropanol solution; 100μL was added to each sample and incubated for 90 minutes at 60°C. The absorbance was measured at 560nm, and a standard curve was used to calculate the hydroxyproline content in each sample.

### Hyaluronan Staining

Hyaluronan was localized as described previously^11^ using a protocol adapted from the Cleveland Clinic’s Program for Excellence in Glycosciences. FFPE tissue sections were rehydrated, equilibrated in PBS, and blocked using avidin and biotin blocking kit (Vector Lab, Burlingame, CA). Normal goat serum was used as a blocking agent and slides were incubated in biotinylated hyaluronan binding protein (HABP, 1:500, CalBiochem, Burlington, MA) for one hour. The Avidin/Biotin complex (Vector Labs) was used to amplify the signal that was then detected with DAB (Vector Labs). Hematoxylin was used as a counterstain. A Leica DMi8 microscope with Leica DFC4500 camera and Las-X software was used to take 20X images (n=3 per mouse). ImageJ was used to measure the area of the submucosa and the percent positive area over a threshold that was kept constant for each image.

### HA Isolation from Murine Vaginal Tissue

HA isolation was performed by biomagnetic precipitation following protocols reported previously with minor modification.^12^ Dynabeads™ M-280 streptavidin-coated superparamagnetic beads (Ref. 11206D, Thermo Fisher Scientific) were first washed according to the manufacturer’s directions and then incubated (2 hours at room temperature under gentle agitation) in 1X PBS with biotinylated versican G1 domain (bVG1, Ref. G-HA02, Echelon Biosciences) at a ratio of 1 µg bVG1 to 100 µg beads. After washing and resuspension in 1X PBS to a concentration of 10 mg/mL, bVG1-conjugated beads were aliquoted into 150 µL volumes. Target tissue was diced and treated with a broad-spectrum protease (proteinase K, Ref. AM2548, Invitrogen, Waltham, MA) following the manufacturer’s directions until no solid material was apparent in the solution. Samples were placed on a heating block at 95°C for 10 min to deactivate protease. The digest was then mixed with an equal volume of phenol:chloroform:isoamyl alcohol, 25:24:1 (Ref. 327111000, Thermo Fisher Scientific, Waltham, MA) and transferred to a phase-lock gel tube (Ref. 2302830, QuantaBio, Beverly, MA) where it was centrifuged (13,000x rpm for 20 min at 20°C) to partition the mixture into an upper aqueous phase containing HA, nucleic acids, and other polysaccharides and a lower organic phase containing digestion proteins and other debris. The process was repeated twice with pure chloroform (Ref. AC423555000, Thermo Fisher Scientific) to remove residual phenol. The resulting solution was added to an aliquot of bVG1 beads and incubated at room temperature for at least 2h for HA capture. Beads were pulled down magnetically from solution and then washed to counter non-specific binding. To elute HA, the beads were incubated in a high salt buffer (6M LiCl, 10 mM Tris, 1 mM EDTA, pH 8.0) for 1h, after which they were again pulled out of suspension by magnetic field and the supernatant was collected for solid-state nanopore (SSNP) analysis.

### Hyaluronan Size Analysis by Solid-State Nanopore Analysis (SSNP)

SSNPs devices (silicon chips supporting a 30 nm thick silicon nitride membrane with a single pore of diameter 7-11 nm) were produced commercially by conventional silicon processing techniques (Norcada, Edmonton, Canada). To perform HA analysis,^13^ a chip was rinsed thoroughly with ethanol and water, dried with filtered air, treated with air plasma (30 W, Harrick Plasma, Ithaca, NY) for 2 min per side, and placed into a custom flow cell fabricated by 3D printing (Carbon, Redwood City, CA). Measurement buffer^14^ (6M LiCl, 10 mM Tris, 1 mM EDTA, pH 8.0) was introduced into independent reservoirs on either side of the SSNP chip and Ag/AgCl electrodes were used connect them to an Axopatch 200B patch-clamp amplifier (Molecular Devices, San Jose, CA). Isolated HA (in measurement buffer from the elution process) was then introduced to one side of the SSNP and measurements were performed by applying 200-300 mV to the opposite side and measuring current at a rate of 200 kHz using a 100 kHz four-pole Bessel filter. Data were collected with a custom LabVIEW program (National Instruments, Austin, TX) through which an additional 5 kHz low-pass filter was applied. Molecular translocations were signified by temporary reductions in the measured current (events), identified using an amplitude threshold of 5σ compared to baseline noise and inclusive only of events with durations in the range of 25 μs - 2.5 ms. The area of each event was mapped to a molecular weight using a calibration curve^15^ produced with a comparable SSNP. Each data set consisted of a minimum of 500 individual events.

### Statistical Analysis

Data was graphed and analyzed using a combination of GraphPad Prism Version 8 and Microsoft Excel. Because our interest was in what effect vaginal injury had at each time point, t-test was used to compare control and bleomycin + epithelial disruption at a single time point. An * was used to designate significance defined as p<0.05. Q-test was used to remove any outlier at 95% confidence.

## RESULTS

Bleomycin is a cellular toxin utilized to induce fibrosis in multiple organ systems in rodent models. Lung fibrosis has been induced using both a low dose bleomycin (1.5U/kg) and high dose bleomycin (2.5U/kg).^15,16^ Similarly, laryngotracheal fibrosis has been induced using bleomycin coated wire brushes which disrupt the epithelial layer.^9,17^ Therefore, to induce vaginal fibrosis, we instilled a low dose (1.5U/kg) and high dose (2.5U/kg) of bleomycin intravaginally. In addition, we also included a high dose bleomycin instillation combined with wire brush epithelial disruption prior to instillation. After 5 exposures over 10 days, and a 3-week healing period, we found neither low nor high dose bleomycin alone induced vaginal fibrosis, evaluated by gene array and trichrome staining (Supplementary Figure 1, Figure 1A). We did however find an increase in pro-fibrotic genes and trichrome staining when high dose bleomycin was combined with epithelial disruption (Supplementary Figure 1). We therefore continued the study using this model.

**Figure 1.**
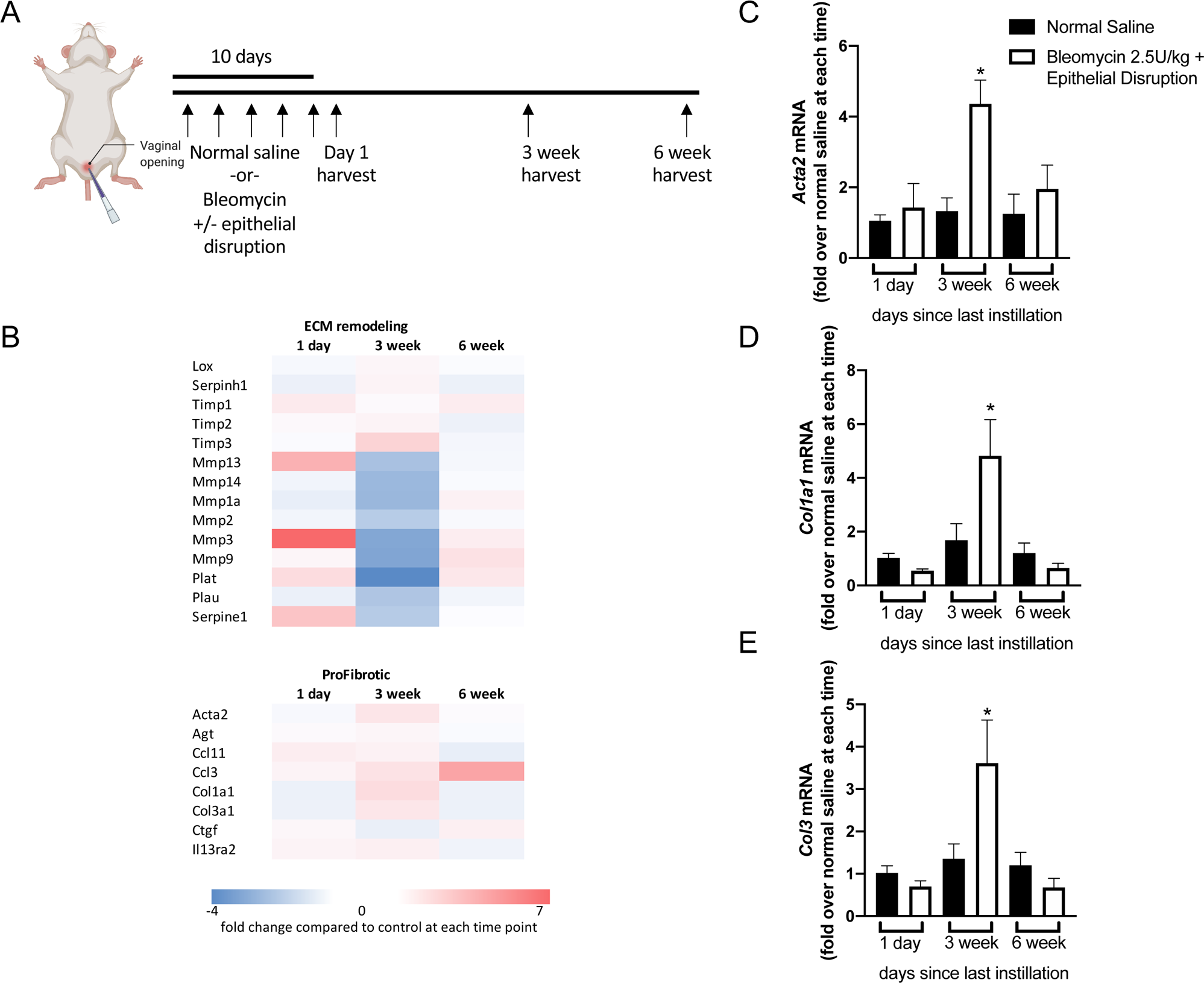
Establishing fibrosis in murine vaginal tissue. (A) Schematized methodology of *in vivo* vaginal instillations where vaginal injury was induced with 2.5U/kg bleomycin (HD) vaginal instillation +/− epithelial disruption. Normal saline vaginal instillation was used as control. Tissue was harvested at 1 day, 3 weeks, or 6 weeks post final instillation. (B) Gene expression array heat map demonstrating major differences of extracellular matrix (ECM) remodelling and profibrotic gene expression using pooled samples from 3 weeks post vaginal injury. No consistent changes are found 1 day or 6 weeks after injury. (C-E) Quantitative PCR confirms gene expression changes observed using the array, with expression of *Acta2*, Col*1a1*, and *Col1a2* being increased in injured vaginal tissue 3 weeks after last instillation compared to control at 3 weeks, but not at 1 day or 6 weeks. (n=4-8, * signifies p≦0.05)

In order to evaluate the progression of vaginal fibrosis, we harvested tissue at three time points after repeated bleomycin plus epithelial disruption, 1 day, 3 weeks, and 6 weeks (Schematic Figure 1A). Using pooled RNA and a fibrosis gene array, we found a decrease in matrix metabolizing genes, particularly those involved in degradation, and an increase in profibrotic and inhibitors of matrix metabolizing genes in bleomycin plus epithelial disruption mice compared to saline control mice 3 weeks after the last instillation. We did not see any consistent trends at 1 day or 6 weeks after bleomycin plus epithelial disruption compared to saline control at the corresponding time point (Figure 1B). This was confirmed by qPCR. We quantified expressions of three pro-fibrotic genes, *Acta2*, *Col1a1*, and *Col3a*, and found that there was no difference between bleomycin + epithelial disruption and saline control at 1 day or 6 weeks after the last instillation. However, all genes were elevated in the bleomycin + epithelial disruption group compared to saline control at 3 weeks after the last instillation (Figure 1C-E). Collectively, this indicates that fibrosis is not present at 1 day, manifests within 3 weeks, and resolves by 6 weeks following the last epithelial disruption and bleomycin intravaginal instillation.

We validated ECM remodeling by analyzing alterations in ECM protein expression by performing trichrome staining to corroborate our gene expression findings. In doing so, the patterns observed in gene expression were verified. There was increased collagen content in mice exposed to bleomycin combined with epithelial disruption compared to saline control after 3 weeks of healing (Figure 2A). We observed no changes at 1 day or 6 weeks after the last vaginal instillation between our experimental mice and control mice (Figure 2A). Hydroxyproline serves as a stabilizer for collagen and is closely linked to collagen levels.^10^ Although not statistically significant, a trend toward elevated hydroxyproline levels was observed in mice exposed to bleomycin and epithelial disruption compared to those administered normal saline, both at 3 weeks (p=0.12) and 6 weeks (p=0.14) after the final instillation (Figure 2B). No changes were seen at 1 day after the last installation between groups (Figure 2B). In conjunction with the gene expression data, these findings provide an insight into the time course of wound healing and fibrosis in our model, where, fibrosis is not established within 24 hours of the final instillation of bleomycin and epithelial disruption. However, by 3 weeks, the presence of fibrosis is evident, and by 6 weeks after the last instillation, fibrosis has mostly resolved.

**Figure 2.**
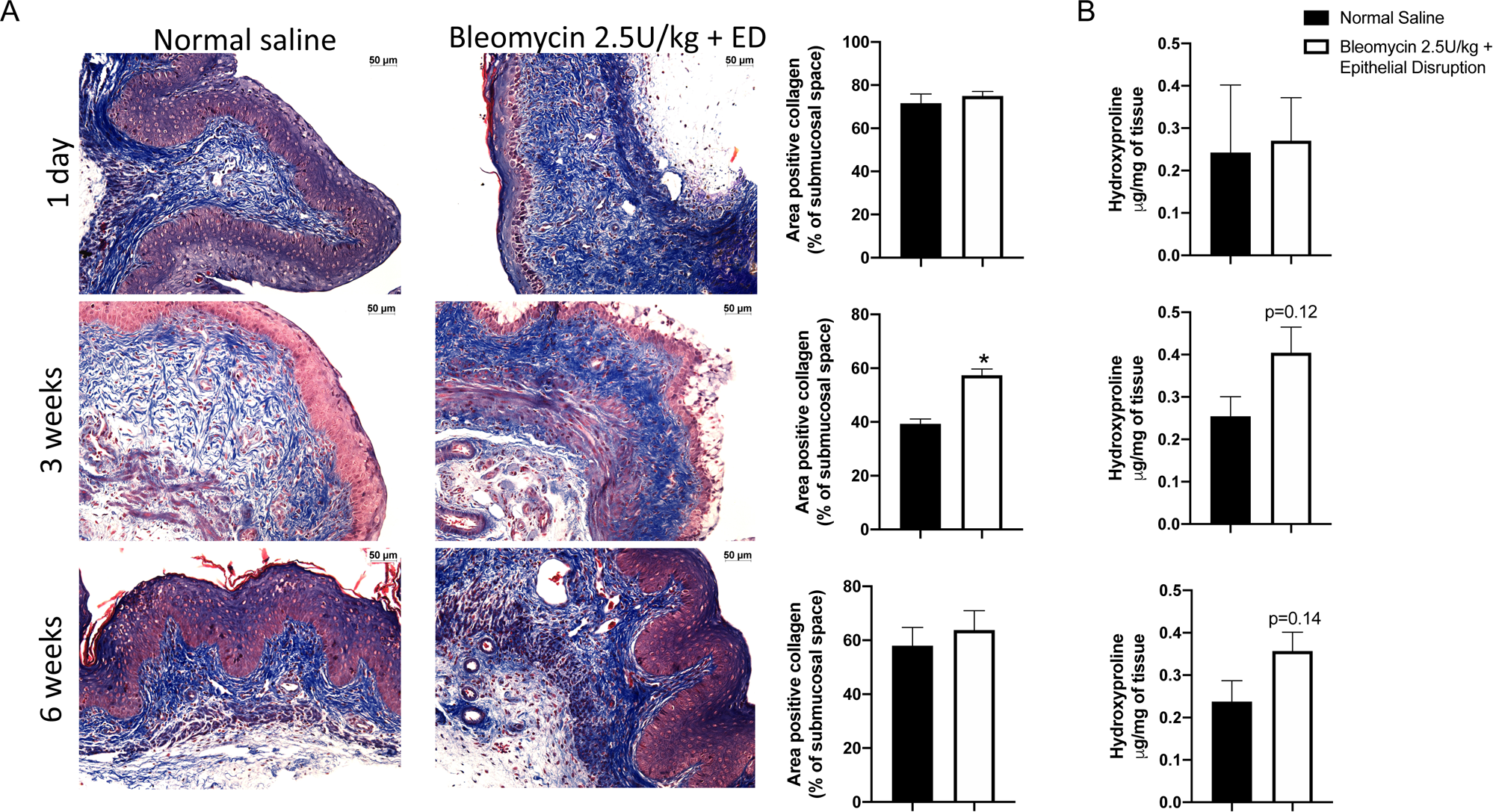
Increased collagen content present at 3 weeks, but not 1 day or 6 weeks post vaginal injury. (A) Representative trichrome images (20X) and the corresponding quantification reveal no differences of collagen content at 1 day or 6 weeks after injury compared to control. However, there is an increased percent positive staining 3 weeks after bleomycin plus ED disruption compared to control. (B) Though not significant, vaginal hydroxyproline content was elevated at 3 weeks and 6 weeks post injury compared to control. (n=4-8, * signifies p≦0.05)

We further conducted an immunohistochemical analysis to investigate vaginal inflammatory cell burden following bleomycin and epithelial disruption injuries. To assess the total leukocyte population (Figure 3), anti-CD45 antibody staining was utilized to stain tissue sections of control and bleomycin plus epithelial disruption mice at day 1, 3 weeks, and 6 weeks post last instillation. Despite vaginal injuries, the total leukocyte population demonstrated a slight but insignificant decrease at all time points. Consequently, during the onset and resolution of vaginal fibrosis, vaginal leukocyte homeostasis was observed, indicating the absence of an active infection or inflammatory response in this model.

**Figure 3.**
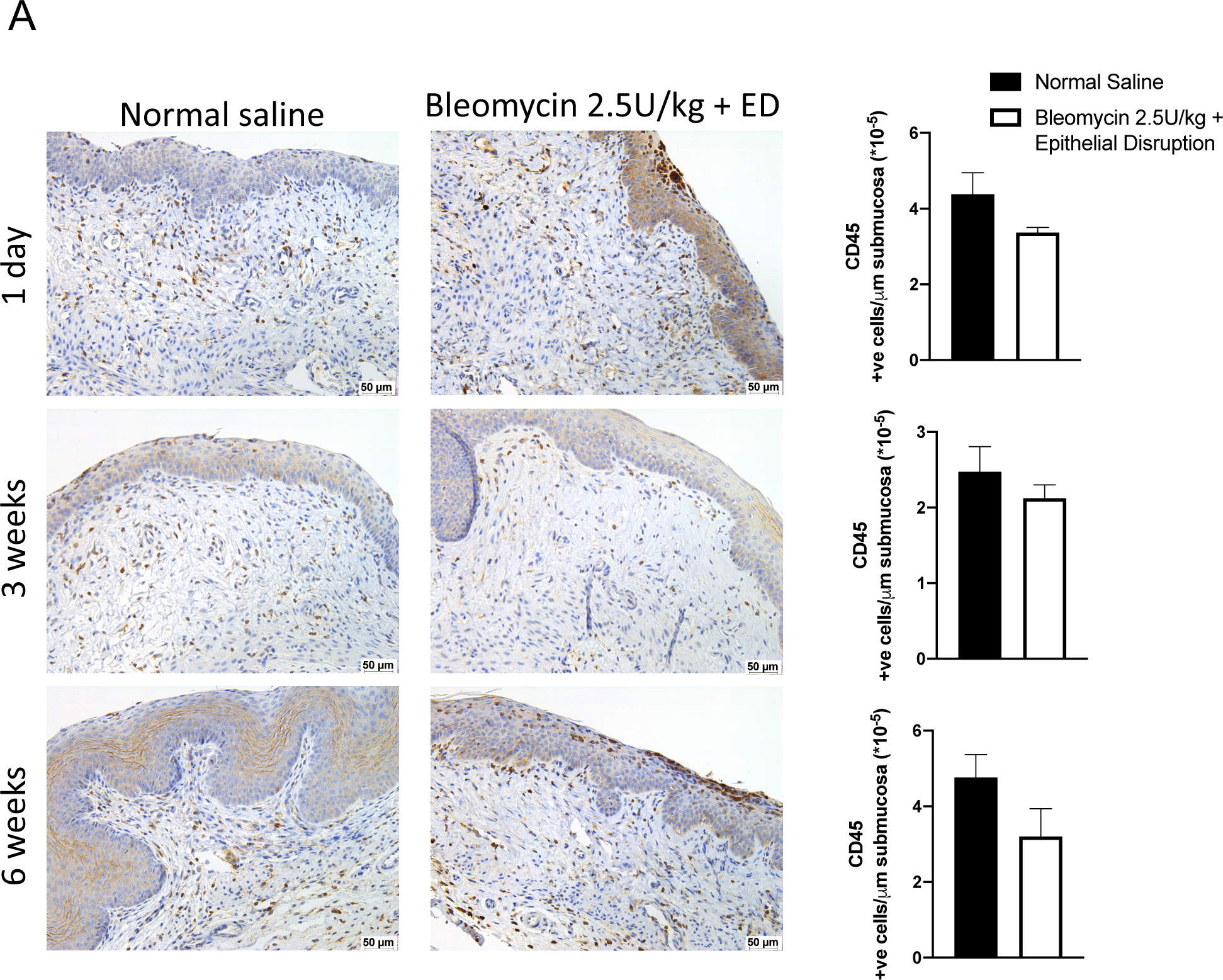
Bleomycin + epithelial disruption does not cause influx of leukocytes. Representative images and quantification of total CD45+ cells demonstrate no significant disruption in basal vaginal inflammatory profile at any time point indicating bleomycin plus epithelial disruption does not have an active inflammatory response.

A dynamic ECM is critical to the outcome of the wound healing process, as it provides not only a scaffold for inflammatory cell entry into the wound bed, but also an active and fluid matrix that directs the healing responses to induce a regenerative or a fibrotic wound healing phenotype. Vaginal fibrosis, which is marked by a significant buildup of collagen in the ECM is noted in the bleomycin with epithelial disruption group at 3 weeks. Nonetheless, the ECM is composed of various other components, including hyaluronan (HA), which is widespread in the vagina.

In order to evaluate vaginal HA response to bleomycin plus epithelial disruption and the development of fibrosis and subsequent resolution, we quantified and characterized the flux of HA molecules within the vagina (Figures 4 and 5). Using HABP to localize HA expression in the wound sections, there appears to be similar HA vaginal content at all time points between control and bleomycin plus epithelial disruption groups (Figure 4A). However, following HA isolation from vaginal tissue, and using an HA ELISA, there is a trend towards decreased HA at 3 weeks (p=0.09) and a significant decrease 6 weeks after the last bleomycin plus epithelial vaginal instillation compared to saline control (Figure 4B). Because HA signaling is closely tied to the molecular size, using HA isolated from the vaginas, we quantified molecular mass by solid-state nanopore analysis. The average HA molecular mass is increased in the experimental mice 3 weeks after bleomycin plus epithelial disruption compared to controls (Figure 5A, B). To understand the size distribution of vaginal HA size, we evaluated the size range of the middle 50% of reads from the solid-state nanopore analysis. Compared to controls, in bleomycin plus epithelial disruption mice there was an increase in this size range at 3 weeks post injury (Figure 5C). No difference was found in either average molecular weight size or size distribution at 1 day or 6 weeks post bleomycin plus epithelial disruption. Collectively, our results demonstrate a reduction in vaginal HA levels following the onset of fibrosis, which remains elevated up to 6 weeks. Moreover, despite the total HA discrepancy, there is a higher molecular weight of HA and a more diverse range of HA sizes within vaginal tissue when fibrosis is established. These observations suggest an irregular HA turnover that persists even after fibrosis has resolved.

**Figure 4.**
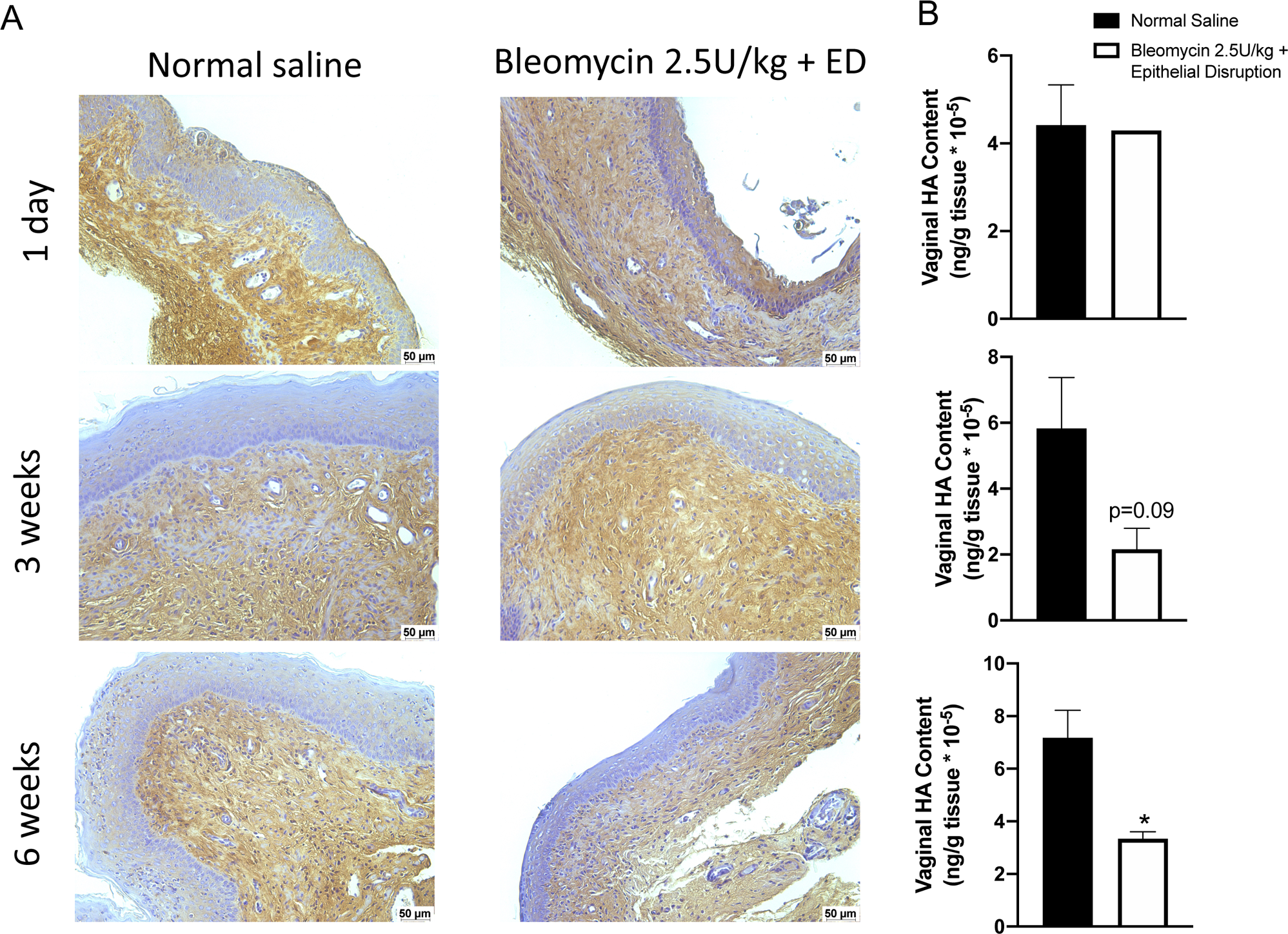
Vaginal hyaluronan (HA) content decreases with fibrosis development and is not recovered by 6 weeks post injury. (A) Representative images of HA presence and distribution in vaginal tissue using HABP 1 day, 3 weeks, and 6 weeks after vaginal injury with bleomycin and epithelial disruption. (B) Quantitative assessment of HA amount in in vaginal tissue by HABP-ELISA shows decrease in total HA content at both 3 weeks and 6 weeks post vaginal injury, but no change 1 day after injury. (n=3, * signifies p≦0.05)

**Figure 5.**
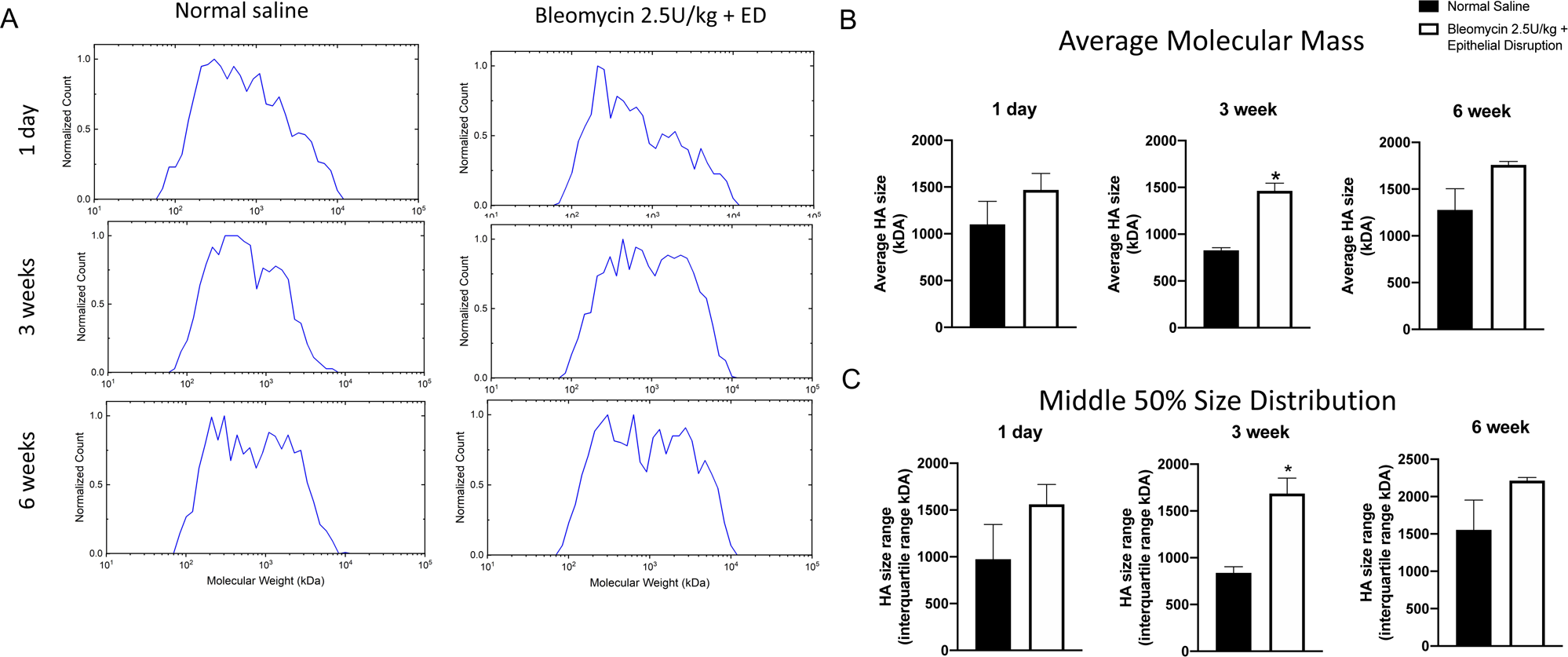
HA molecular weight size distribution is altered following bleomycin plus epithelial disruption vaginal injury. (A) Representative histograms of solid state nanopore analysis to measure HA size in vaginal tissue. (B) Quantification of individual reads showed average HA molecular mass is increased at all time points in bleomycin + ED mice compared to controls and statistically significant at 3 weeks. (C) The average size distribution over middle 50% of reads in bleomycin + ED mice compared to controls is larger 3 weeks after injury, while no difference is found 1 day or 6 weeks after vaginal injury. This suggests wider range of size of HA at 3 weeks post injury compared to control. (n=3, * signifies p≦0.05)

## DISCUSSION

In this work, we have established and characterized a novel, reproducible, and site specific murine vaginal fibrosis model. Moreover, we have demonstrated the time course of vaginal wound healing from injury to fibrotic state, and subsequent resolution. This novel murine model eliminates many of the issues of previous murine vaginal injury models including small injury site, accelerated murine vaginal wound healing kinetics, and difficulty in creating reproducible surgical injuries leading to fibrosis while recapitulating clinical relevance using a gynecologic curettage technique.^3^ This murine model will allow the development of novel preventions and/or treatments for vaginal fibrosis, a debilitating condition which plagues millions of women worldwide. Additionally, this model will serve as a tool to study the cellular and molecular mechanisms governing the spectrum of mucosal wound healing dynamics using the vagina as an archetype of these responses to injury.

With this model, we have also characterized several important alterations at the genetic, cellular, and tissue levels related to the vaginal wound healing spectrum from fibrosis to regeneration. First, we clarified the genetic markers for vaginal states characterized by fibrosis and regeneration. In line with other tissues, our investigation revealed elevations in genes that promote fibrosis such as *Acta2*, *Col1a1*, and *Col3*, as well as changes in genes that regulate the breakdown and renewal of extracellular matrix (ECM) components. The changes included overall reductions in matrix metalloprotease enzymes and increases in tissue metalloprotease inhibitors. These findings support the notion that the process of fibrosis involves a coordinated increase in ECM collagen production and a decrease in ECM degradation. The gene changes translated to vaginal fibrosis at 3 weeks after injury, and were confirmed both histologically and molecularly. We also demonstrate the need for epithelial disruption combined with chemical injury to be vital in inducing vaginal fibrosis, as chemical injury alone did not result in vaginal fibrosis.

Importantly, this model also contributes to the paradigm that the ECM is not a passive scaffold for wounds but actively participates in progression of both regeneration and fibrosis. It is possible that the vaginal ECM can repair itself following some degree of harm, such as with the use of bleomycin alone. However, once a certain level of damage is surpassed, the genetic mechanisms that lead to fibrosis are triggered, making it inevitable unless an intervention is implemented. Furthermore, our findings demonstrate that fibrosis has been fully resolved within six weeks of injury, showing the murine vagina possesses an impressive capability to restore its equilibrium and regain homeostasis.

Inflammation often proceeds tissue injury to repair the tissue. Chronic inflammation has long been thought to be a contributing factor for fibrosis,^18^ however recent work has suggested that inflammation, macrophages specifically, are necessary to counteract the fibrotic buildup.^19^ The vagina is unique in that it has a sustained homeostatic leukocyte population necessary to protect the tissue from potential infection.^20^ After inducing vaginal injury through repeated exposure to bleomycin in conjunction with epithelial disruptions, we observed that the total leukocyte populations remained relatively constant throughout the course of injury and regeneration, with no significant changes. As fibrosis progressed, the leukocyte population demonstrated a slight but not significant reduction. Notably, even six weeks after injury, the leukocyte population remained slightly lower compared to control mice, despite the excess collagen in vaginal tissue having been resolved.

Hyaluronan (HA) is one of the predominant ECM components in the vaginal tissue.^21^ This highly bioactive glycosaminoglycan is distributed widely in vaginal stroma and surrounding epithelial cells.^22–24^ HA is a major hallmark in tissue injury with sudden increase and turnover in the ECM.^25^ Differences in HA synthesis, accumulation, and degradation patterns can lead to variable wound healing outcomes upon injury.^26^ The effect HA has on healing is largely dependent on its molecular weight. Low molecular weight HA (LMW-HA) promotes a pro-inflammatory environment, whereas high molecular weight (HMW-HA) dampens pro-inflammatory effects.^27–29^ The balance of LMW-HA to HMW-HA can also modulate inflammation during wound healing. In fact, the wound repair process leading either to fibrosis or regeneration can be understood as a disordered or coordinated HA balance both in size and amount.^30^ Regenerative tissue such as fetal skin, has a high abundance of HMW-HA and tissues prone to scaring, i.e., aged skin, has lower amounts of HMW-HA.^26,31^ Although HA is a key player in driving wound healing and fibrotic outcomes in other tissues, its specific role in mediating mucosal vaginal tissue responses to **i**njury remains uncharacterized. As such, we have begun to characterize the flux of HA molecules to the vaginal wound bed both by total amount and molecular mass. Total HA amounts decreased during collagen accumulation; however, the average molecular mass was highest in the vaginal tissue at 3 weeks when fibrosis is present and it remains elevated at 6 weeks, despite the resolution of fibrosis. Additionally, when we consider the size range of HA, it is wider at the time of vaginal fibrosis. We have demonstrated therefore that HA turnover is altered during vaginal fibrosis progression, but re-establishing homeostasis of HA lags collagen in the injury-repair progression from injury to fibrosis to regeneration.

The findings from this study indicate that vaginal fibrosis can be induced by epithelial disruption in conjunction with bleomycin vaginal instillation, leading to a noticeable increase in collagen synthesis and decrease in ECM degradation, ultimately resulting in established fibrosis within three weeks of the injury regimen. Additionally, we observed changes in other ECM components, specifically hyaluronan (HA), in this injury model. Our findings revealed that the murine vagina possesses impressive regenerative abilities, able to fully recover within six weeks. These results provide a valuable platform for further elucidation of the mechanisms that underpin vaginal fibrosis, as well as potential preventive and treatment strategies for the many women who suffer from this condition.

One of the limitations of this study is that the estrus cycles in the mice were not in sync. Furthermore, the signaling effects of estrogen on this injury model were not studied. Fundamentally, estrogen is known to play a key role with respect to inflammatory burden modulation and angiogenic direction of both dermal and vaginal wound healing.^3^ Therefore, estrogen’s specific and intertwined role in mediating collagen synthesis and HA responses to vaginal tissue injury will be exciting future directions to interrogate. Additionally, though bleomycin is a toxic anti-cancer agent known to cause fibrosis in other mucosal organs,^9^ we recognize its mechanism of action leading to fibrosis is likely different than surgically related or radiation induced vaginal fibrotic outcomes. Lastly, we recognized the potential small sample size used for HA analysis, and further studies to increase power are planned.

Overall, understanding the fundamental mechanisms that govern how the vaginal mucosa heals, both pathologically and regeneratively, will allow new insights into the design of personalized therapies to improve patient outcomes. With novel mouse models such as this bleomycin combined with epithelial disruption induced fibrosis model, we can more robustly define the markers of pro-fibrotic phenotypes in mucosal tissues. Additionally, due to the ability of the murine vagina to resolve the established fibrosis, we can harness this model at different time points to understand both the progression and resolution of fibrosis. Future directions include elucidating the interactions between tissue stretch and estrogen regulation of cellular processes that underpin the spectrum of vaginal wound repair. Whether pathologic states of disordered wound healing could be susceptible to regulation by estrogen and biomechanical stretch and shift towards a regenerative phenotype sooner may promise solutions for women undergoing curettage or vaginal surgical repairs, and vaginal radiation therapies.

## KEY FINDINGS

- A murine vaginal injury model is developed to provide a reproducible and clinically relevant tool for studying vaginal fibrosis and investigating cellular and molecular mechanisms involved in mucosal wound healing dynamics.
- Both low (1.5U/kg) and high doses (2.5U/kg) of bleomycin alone were insufficient to induce vaginal fibrosis. However, when combined with epithelial disruption using a wire brush, high dose bleomycin resulted in increased pro-fibrotic gene expression and collagen deposition, indicating the induction of vaginal fibrosis.
- Vaginal fibrosis manifested within three weeks and resolved by six weeks after the last instillation of bleomycin and epithelial disruption.
- Vaginal leukocyte populations remained relatively constant throughout the injury and regeneration process, suggesting the absence of an active infection or inflammatory response during vaginal fibrosis.

## ACKNOWLEDGEMENTS AND FUNDING SOURCES

This work was graciously supported by NIGMS 1K08GM135638-01. Additionally, the authors acknowledge the support of the entire Laboratory for Regenerative Tissue Repair team at the Texas Children’s Hospital.

## AUTHORS CONTRIBUTION

**Jennifer McCracken**: conceptualization (supporting); writing-original draft (supporting); writing-review and editing (supporting); data curation (equal); formal analysis (supporting); investigation (supporting). **Gisele Calderon:** conceptualization (supporting); writing-original draft (supporting); writing-review and editing (supporting); data curation (equal); formal analysis (supporting); investigation (supporting). **Lishore Kumar:** writing-review and editing (lead); formal analysis (supporting). **Swathi Balaji:** conceptualization (supporting); writing-original draft (supporting); writing-review and editing (supporting); data curation (equal); formal analysis (supporting); investigation (supporting). **Julie Hakim:** conceptualization (lead); writing-original draft (lead); formal analysis (lead); supervision (lead); project administration (lead); investigation (equal), funding acquisition (lead).

## AUTHOR DISCLOSURE AND GHOSTWRITING

The authors of this manuscript, titled “Unveiling Vaginal Fibrosis: A Novel Murine Model Using Bleomycin and Epithelial Disruption”, declare no conflicts of interest that could be perceived as potentially influencing the objectivity and impartiality of the reported research.

## ABOUT THE AUTHORS

**Jennifer McCracken, PhD** was a research associate at Baylor College of Medicine in OBGYN and is currently a student at University of Kansas School of Pharmacy.

**Gisele A. Calderon, PhD** was a research associate at Baylor College of Medicine in OBGYN and is currently a Scientist, R&D, Regenerative and Biological Exploration at Integra Lifesciences.

**Lishore Kumar** is a high schooler at Tomball Memorial High School pursuing research at Baylor College of Medicine.

**Swathi Balaji, PhD** is an assistant professor in the Department of Surgery at Baylor College of Medicine. She has been a member of the Wound Healing Society.

**Felipe Rivas, PhD** was a research associate at Wake Forest University in Biomedical Engineering and is currently a R&D engineer at Charter Medical LLC.

**Dorothea Erxleben** is a PhD student at Wake Forest University in the School of Biomedical Engineering and Sciences.

Adam Hall, PhD is an associate professor in the School of Biomedical Engineering at Wake Forest University School of Medicine.

**Julie CE Hakim, MD, FRCS(C), FACOG** Dr. Julie Hakim is a Canadian and U.S. board-certified OBGYN and Assistant Professor in OBGYN at Baylor College of Medicine and Pediatric Gynecology at Texas Children’s Hospital in Houston, Texas.

## ABBREVIATIONS AND ACRONYMS

ECM: Extra cellular matrix
ED: Epithelial disruption
HA: Hyaluronan
HABP: Hyaluronan binding protein
HMW-HA: High molecular weight hyaluronan
LD: Low dose
LMW-HA: Low molecular weight hyaluronan
SSNP: Solid state nanopore

---

**Supplemental Figure 1.**
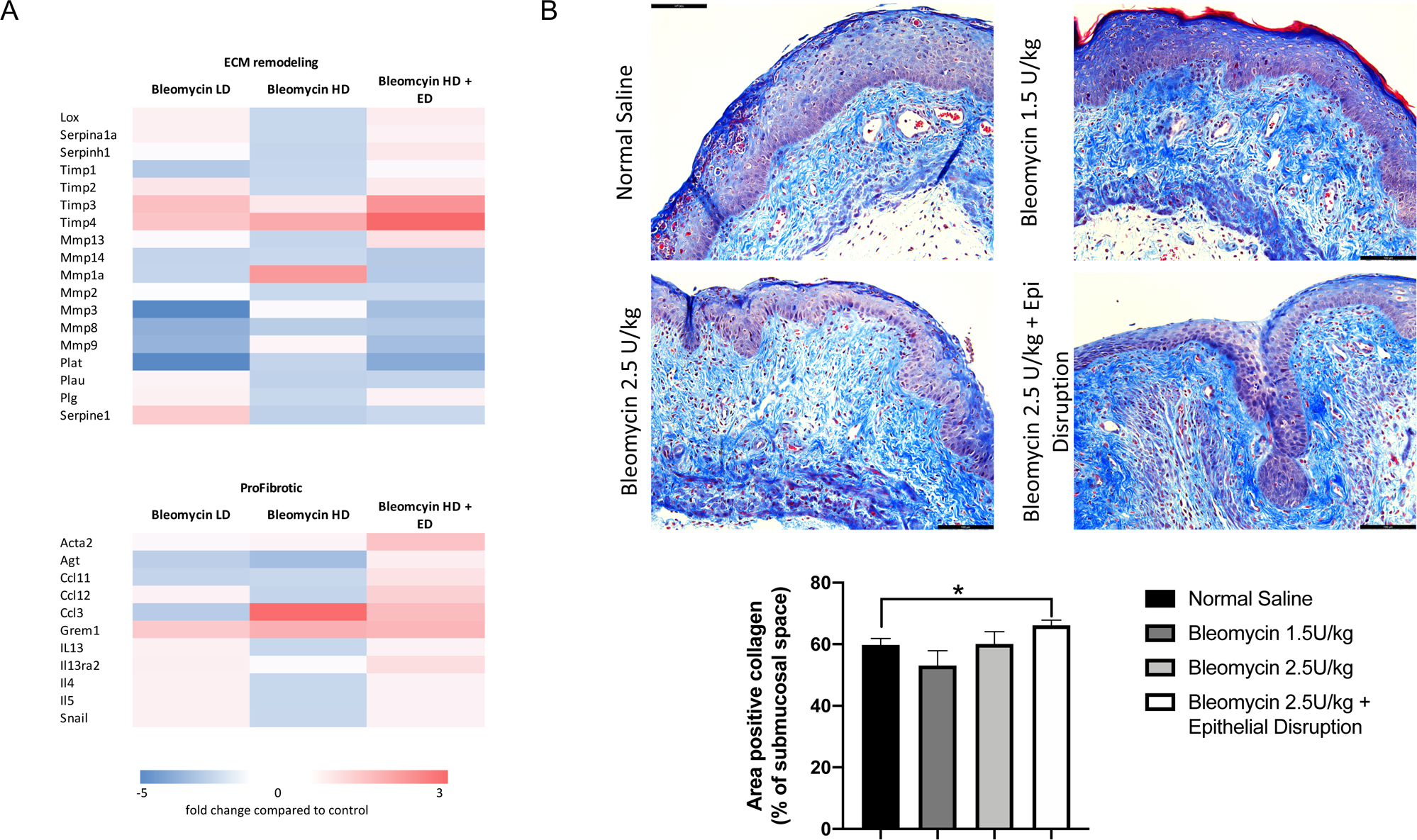
Establishment of fibrotic wound healing model in murine vaginal tissue 3 weeks after bleomycin with or without epithelial disruption (ED) exposure. (A) Gene expression array heat map demonstrating consistent differences in ECM remodeling and profibrotic gene expression in bleomycin high dose (HD) plus ED mice compared to control, particularly increased pro-fibrotic genes. No expression pattern is observed in low dose (LD) bleomycin or high dose (HD) bleomycin alone compared to control. (B) Representative 20X trichrome images and quantification of control, LD and HD bleomycin, and HD bleomycin plus ED demonstrating significant differences in collagen content between control and HD bleomycin plus ED. (scale bar = 100μm, n=7-8, * signifies p≦0.05)

